# The oocyte zinc transporter *Slc39a10/Zip10* is a regulator of zinc sparks during fertilization in mice

**DOI:** 10.1101/2024.09.11.612381

**Authors:** Atsuko Kageyama, Narumi Ogonuki, Takuya Wakai, Takafumi Namiki, Yui Kawata, Manabu Ozawa, Yasuhiro Yamada, Toshiyuki Fukada, Atsuo Ogura, Rafael A. Fissore, Naomi Kashiwazaki, Junya Ito

**Author notes:** Department of Life Science Frontiers, Center for iPS Cell Research and Application (CiRA), Kyoto University, Kyoto, 606-8507, Japan (Takafumi Namiki). Correspondence: Junya Ito, Laboratory of Animal Reproduction, Graduate School of Veterinary Sciences, Azabu University, Sagamihara, 252-5201, Japan.

## Abstract

In all vertebrates studied to date, a rise(s) in intracellular calcium is indispensable for successful fertilization and further embryonic development. Recent studies demonstrated that zinc is ejected to the extracellular milieu, the ’zinc spark’, and follows the first few calcium rises of fertilization. However, the role of the zinc sparks in fertilization and development, and the supporting influx mechanism(s) are unknown. In this study, we focused on zinc transporters *Zip10/Slc39a10* which was expressed in mouse oocytes through follicular development, and investigated the oocyte-specific deficient mice for *Zip10* (*Zip10^d/d^*: *Zip10^flox/flox^ Gdf9^Cre/+^*). *Zip10* mRNA or ZIP10 protein was expressed throughout folliculogenesis in the oocyte or plasma membrane, respectively. The number of ovulated oocytes was examined in *Zip10^d/d^* mice, and no change from the number of oocytes was observed. *Zip10^d/d^* oocytes decreased zinc level in the oocytes, but did not affect maturation and metaphase II spindles formation. Fertilization-induced calcium oscillations were present in *Zip10^d/d^* oocytes, but zinc sparks were not observed. Despite other events of egg activation proceeding normally in *Zip10^d/d^*oocytes, embryo development into 4-cells and beyond was compromised. We show here for the first time that the zinc transporter ZIP10 contributes to zinc homeostasis in oocytes and embryos, highlighting the role of labile zinc ions in early development.

**Submission information:** The zinc transporter, *Slc39a10/Zip10*, is required for the zinc sparks of fertilization in mice.

## Introduction

In mammalian fertilization, the sperm factor is released into the ooplasm to activate the oocyte (Yoon and Fissore, 2007). It is now believed that phospholipase C zeta (PLCζ) (Yoon and Fissore, 2007; Saunders et al., 2002) is the sperm factor and it induces in the ooplasm repetitive increases in the intracellular Ca^2+^, termed ’calcium oscillations.’ These oscillations play an essential role in triggering the oocyte activation events, such as cortical granule exocytosis, the block of polyspermy, meiotic resumption and exit from MII arrest (Xu et al., 1994; Jones, 2005; Ducibella et al., 2006; Ito et al., 2011; Sugita et al., 2024). A calcium increase is highly conserved in many species as the trigger of egg activation and is vital for successful fertilization.

In recent years, zinc ions as well as calcium ions, are thought to play important roles during fertilization (Suzuki et al., 2010; Allouche-Fitoussi and Breitbart, 2020; Kageyama et al., 2022). Zinc ion is an essential trace element and the second most abundant transition ion in the human body after iron (Lubna and Ahmad, 2023). Zinc homeostasis is essential for optimal metabolic function in the reproductive process in mammals (Pascua et al, 2020), therefore zinc deficiency causes abnormalities such as fetal teratogenicity, long gestation periods, problematic labor, low birth weight and weak offspring (Favier, 1992; Bedwal and Bahuguna, 1994). Zinc also has important role in the male and female germ cells (Allouche-Fitoussi and Breitbart, 2020; Kageyama et al., 2022). Especially in the mouse female germ cells, zinc-deficient condition caused profound defects in oocyte maturation, cumulus cell expansion and ovulation cycle (Tian and Diaz, 2012). Several studies have been conducted in mice to understand how zinc regulates oocyte maturation. Zinc was reported to be a critical regulator of meiosis throughout oocyte maturation, including maintaining release from the first and second meiotic arrest (Kong et al., 2012). In addition, zinc levels of oocytes were found to increase by 50% as the oocyte progressed through meiosis (Kim et al., 2010; Bernhardt et al., 2011).

In 2011, Kim *et al*. reported the importance of zinc during mammalian fertilization (Kim et al., 2011). Zinc is predominantly stored in vesicles that are symmetrically arranged along the oocyte cortex at the GV stage, and zinc-stored vesicles (cortical granules) are located away from the spindle and form a hemispherical pattern at the MII stage (Kim et al., 2011; Que et al., 2015; Kong et al., 2014; Que et al., 2019; Jo et al., 2019; Que et al., 2017). After the penetration of sperm, it has been induced a ‘zinc spark’ that releases billions of labile zinc ions from the cortical granules, and the total zinc content of the oocytes decreases by 10-20% in at the end of fertilization in mice (Kim et al., 2011; Lee et al., 2020). In addition, these events immediately follow a series of calcium oscillations (Kim et al., 2011). Zinc spark has also been observed in the cattle, non-human primate, human and Xenopus laevis oocytes (Kim et al., 2011; Que et al., 2019; Duncan et al., 2016; Seeler et al., 2021), suggesting that this phenomenon is a highly conserved event in vertebrate. More surprisingly, full-term development of mouse embryos has been reported by chelation of Zn^2+^ ions without Ca^2+^ release, suggesting the depletion of zinc ions in oocytes may be sufficient for oocyte activation(Suzuki et al., 2010). Zinc has also been reported to impact the sperm during fertilization in mice. Zinc accumulation in the zona pellucida increases fibril binding along the glycoprotein matrix and decreases the number of sperm that can reach the fertilized oocytes (Que et al., 2017; Aonuma et al., 1981; Tokuhiro and Dean, 2018). After the zinc sparks, the released zinc affects the forward motility of sperm to prevent their passage through the zona matrix (Tokuhiro and Dean, 2018). Zinc spark is associated with the release of ovastacin (*Astl*, the official gene name), which is required for the zona reaction, and functions as a polyspermy block mechanism that is initiated only a few minutes after fertilization (Tokuhiro and Dean, 2018).

The transport of zinc ions into and out of cells is regulated by zinc transporters. In mammals, 14 zinc ion importers, called as ZIP (ZIP1-ZIP14), have been identified (Fukada and Kambe, 2011; Takagishi et al., 2017; Kambe et al., 2015). Previous study showed that mammalian oocytes regulate zinc uptake through two maternally derived and cortically distributed zinc transporters, ZIP6 and ZIP10 (Kong et al., 2014). Further, they reported that targeted disruption using *Zip6*-and *Zip10*-specific morpholino injection or antibody incubation induced alteration of the intracellular labile zinc content, spontaneous resumption of meiosis from the PI arrest and premature arrest at a telophase I-like state (Kong et al., 2014). It is clear from these reports that ZIP6 and ZIP10 are involved in zinc transport in oocytes, but the function is not elucidated.

In this study, we generated oocyte specific *Zip6* and *Zip10* conditional knockout mice and examined the function of ZIP10 in the oocytes and the importance of zinc homeostasis during fertilization and embryonic development. This study provides clues that elucidate its role in fertilization and embryonic development, which is still largely unknown. In addition, this is the first report confirming the function of the zinc transporter in oocytes, which will contribute to future research.

## Results

### Zinc transporters, ZIP6, and ZIP10, are expressed in mouse oocytes through follicular development

First, using *in situ* hybridization and immunofluorescent staining, we examined the expression of ZIP10 during follicular development. As shown in Fig. 1A, *Zip10* mRNA was expressed from the primordial oocyte (Arrow). It continued to be expressed in oocytes of primary, secondary, and antral follicles. ZIP10 protein was also expressed in the plasma membrane of primordial oocytes (Fig.1B; Arrow). We also confirmed the expression of ZIP10 protein at the plasma membrane of oocytes of primary, secondary, and antral follicles. Although ZIP6 was also expressed in oocytes throughout folliculogenesis (Fig.1C; Arrow), it displayed nuclear localization in oocytes and granulosa cells of primary, secondary, and antral follicles. Fig. 1D showed zona pellucida and granulosa cells through the follicular development.

**Fig. 1.**
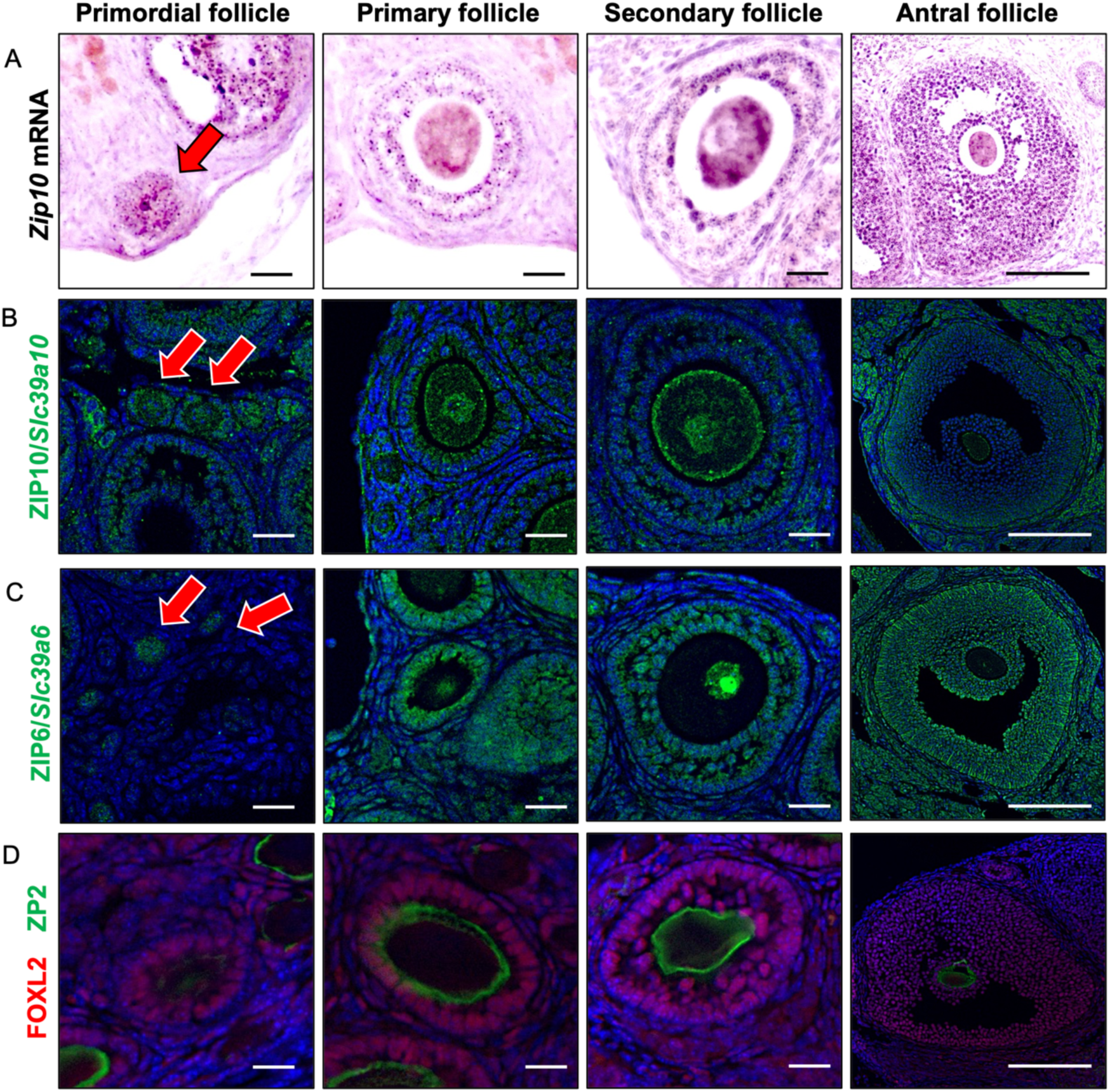
Expression of ZIP6 and ZIP10 in mouse ovary. **(A**) In situ hybridization in the mouse ovary showed ZIP10 expression in oocyte and granulosa cell from primordial, primary, secondary and antral follicle. Arrow indicates primordial follicular oocyte. (**B**) Immunofluorescent staining for ZIP10 (green) in the mouse ovary showed ZIP10 expression in oocyte membrane. Arrow indicates primordial follicular oocyte. (**C**) Immunofluorescent staining for ZIP6 (green) in the mouse ovary showed ZIP6 expression in oocyte nucleus and granulosa cells. Arrow indicates primordial follicular oocyte. (**D**) Immunofluorescent staining showed ZP2 (green; zona pellucida) and FOXL2 (red; granulosa cells) in the mouse ovary. It was observed that ZP2 was not present in the primordial follicle; however, it was present in the primary, secondary and antral follicles. Furthermore, FOXL2 was observed at granulosa cells of all stage follicles. Scale bar: 20 µm (primordial, primary and secondary follicle); 150 µm (antral follicle) (**A-D**).

### Phenotype of oocyte-specific *Zip6^d/d^* and *Zip10^d/d^*female mice

To elucidate the roles of *Zip6* and *Zip10* in the mouse oocytes, oocyte-specific *Zip6* (*Zip6^d/d^*) and *Zip10* (*Zip10^d/d^*) knockout mice were generated (Fig. S1A-C). We examined whether ZIP6 and ZIP10 protein expression was absent in *Zip6^d/d^* and *Zip10^d/d^* oocytes (Fig. S1D). *Zip6^f/f^* and *Zip10^f/f^*mice were used as controls, respectively. We examined the number of ovulated oocytes or defects in oocyte maturation. After superovulation, the number of ovulated oocytes collected from the oviduct in *Zip10^d/d^*mice was 22.1 ± 2.5 oocytes/mouse (Fig. 2A), which was equivalent to that in *Zip10^f/f^* mice (18.7 ± 2.9 oocytes/mouse, *p* > 0.05). The numbers of ovulated oocytes in *Zip6^d/d^* and *Zip6^f/f^*mice were also equivalent (28.7 ± 3.6 and 32.8 ± 3.3 oocytes/mice, respectively, *p* > 0.05) (Fig. S2A). As for oocyte maturation, the rate of oocytes with a first polar body at 10h, 12h, and 14h in the *Zip10^d/d^* group was not different than that of *Zip10^f/f^* group (*Zip10^d/d^*: 58.7%, 61.9%, and 79.4%, *Zip10^f/f^*: 50.8%, 54.1% and 70.5%, respectively, *p* > 0.05, Fig. 2B). In addition, *Zip10^d/d^* oocytes display intact metaphase spindles just as in the *Zip10^f/f^* groups (Fig. 2C).

**Fig. 2.**
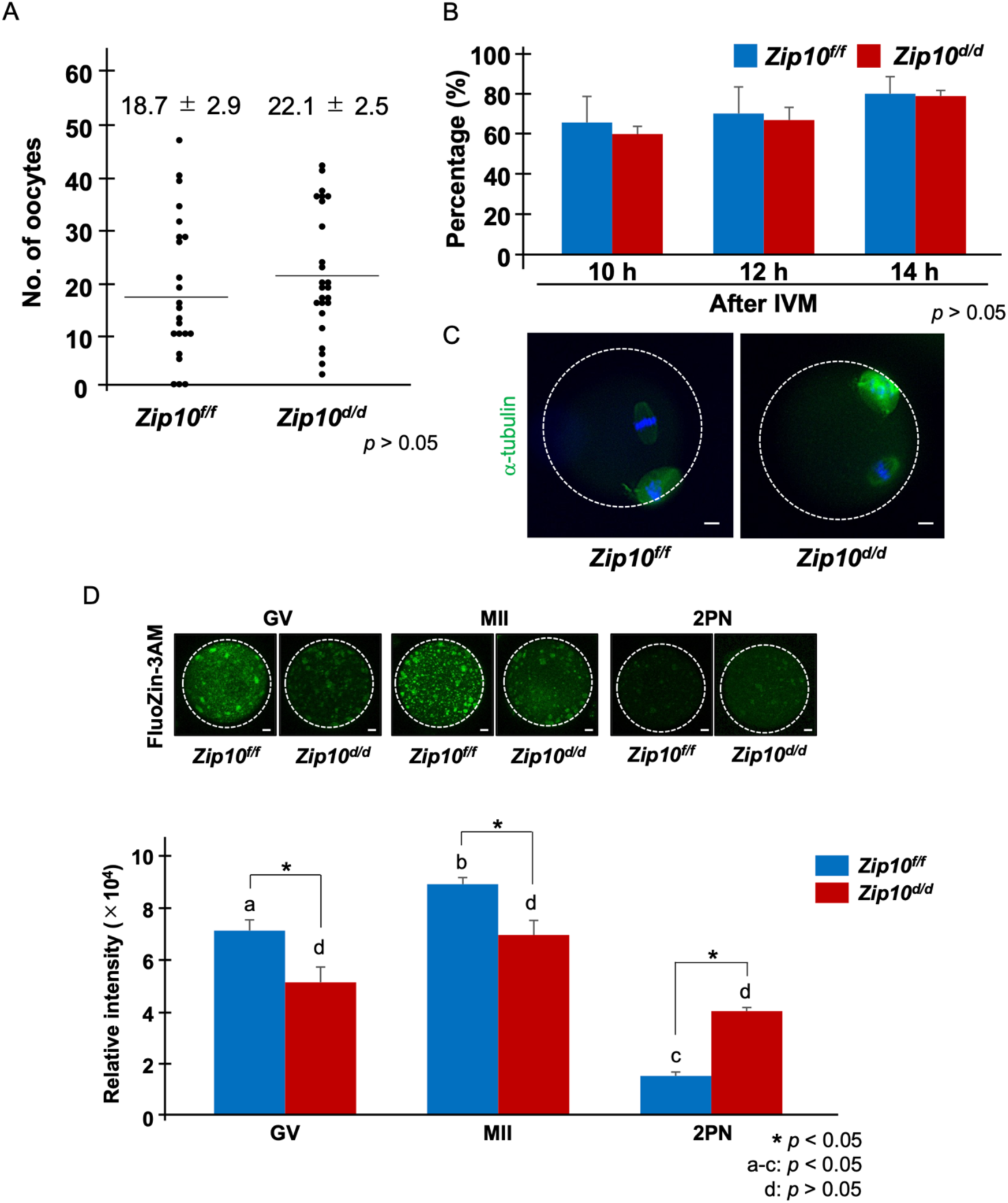
Number of collected oocytes and dynamics of labile zinc ion in *Zip10^d/d^* mice. (**A**) The results of average number of oocytes in each group. Data represents the average ± SEM. These experiments were repeated at least thrice. Statistical differences were calculated according to student’s *t-*test (*p* > 0.05; no significant difference). (**B**) The percentage of extrusion of first polar body at 10, 12 and 14 h after IVM. These experiments were repeated at least thrice. Statistical differences were calculated according to student’s *t*-test (*p* > 0.05; no significant difference). (**C**) The morphology of spindle and chromosome organization in *Zip10^f/f^* and *Zip10^d/d^* MII oocytes at 14 h after IVM. Anti–α-tubulin antibody (green) was used to stain the spindles. Chromosomes were stained with DAPI (blue). The scale bar represents 10 μm. (**D**) Comparison with the fluorescence intensity of intracellular labile zinc ion in GV, MII and 2PN. The upper images showed the fluorescence of intracellular labile zinc ion of oocyte or embryo treated with 2 µM FluoZin-3AM for 1h. Representative images are shown. The white dotted circles indicate the positions of oocytes and embryos. Scale bars denote 10 μm. The lower part showed the fluorescence intensity of labile zinc ions in oocytes or embryos. Data represent the average ± SE of the experiments. For each experiment, 10–20 oocytes/embryos were stained and used for the measurement in each stage of the experiment, and these experiments were repeated three times. Statistical differences were calculated according to the welch’s *t*-test. Different letters represent significant differences (p < 0.05).

Next, we examined the levels of labile zinc in germinal vesicle (GV), MII, and two pronuclei (2PN) zygotes by comparing the fluorescence intensity following loading with the dye, FluoZin-3AM (Fig. 2D). The FluoZin-3AM fluorescence intensity in GV and MII oocytes of the *Zip10^d/d^* group was lower than the *Zip10^f/f^* group (*p* < 0.05). After fertilization, the fluorescence intensity in *Zip10^f/f^* zygotes decreased dramatically, and also decreased for the *Zip10^d/d^* group, although it remained higher than for *Zip10^f/f^*oocytes. When compared within group, the fluorescence intensity in *Zip10^f/f^*between GV and MII oocytes was significantly different (GV vs. MII and MII vs 2PN; *p* < 0.05, respectively), but the stages of *Zip10^d/d^* oocytes were not different despite clear trends (*p* > 0.05). The levels of zinc fluorescence intensity in the *Zip6^d/d^* group were not different from the controls *Zip6^f/f^*, which decreased markedly at fertilization (Fig. S2B; *p* > 0.05).

### The zinc sparks of *Zip10^d/d^* oocytes were suppressed after fertilization or artificial activation

To determine if the absence of ZIP6 and ZIP10 influenced the detection of Zn^2+^ sparks associated with fertilization, we monitored extracellular zinc and intracellular calcium during fertilization of *Zip6^d/d^* and *Zip10^d/d^* oocytes (Fig. S3 and 3A). *Zip10^f/f^* oocytes displayed the expected calcium oscillations following fertilization, and a zinc spark followed the first calcium rise (Fig. 3A upper side, movies 1 and 2). In contrast, *Zip10^d/d^*oocytes did not release zinc ions immediately after the first calcium spike, despite mounting normal calcium oscillations (Fig. 3A lower side, movies 3 and 4). In *Zip6^d/d^*oocytes, a zinc spark occurred immediately after the first intracellular calcium rise at fertilization, just as in control *Zip6^f/f^* oocytes (Fig. S3, movie S1-S4). The extracellular zinc sparks were examined following artificial oocyte activation of mouse oocytes with ionomycin. The zinc sparks occurred immediately after the intracellular calcium rise in *Zip10^f/f^*oocytes (Fig. 3B upper side, movies 5 and 6), however, *Zip10^d/d^* oocytes did not release zinc ions after the calcium spike (Fig. 3B lower side, movies 7 and 8).

**Fig. 3.**
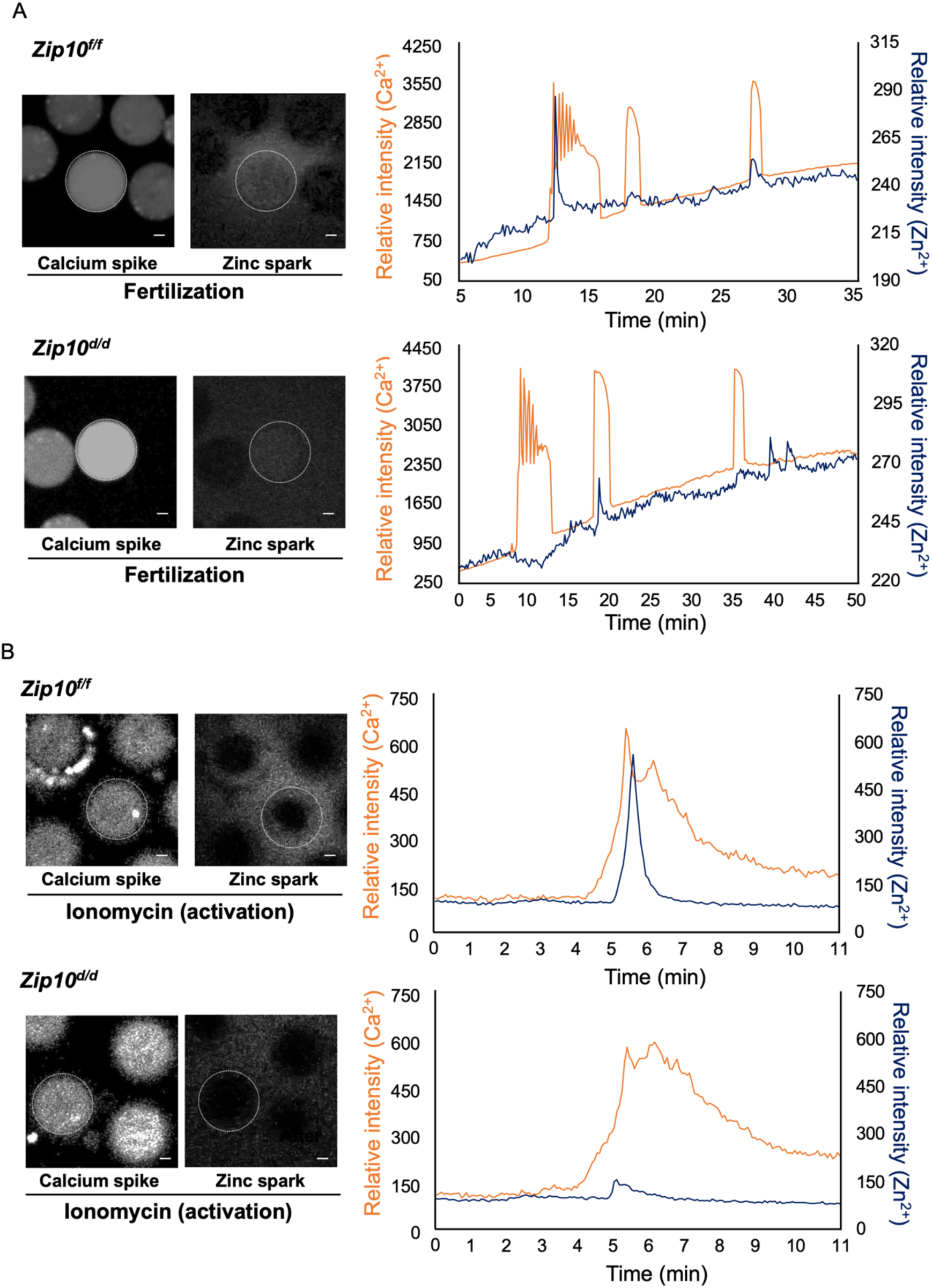
Measurement of calcium spike and zinc spark in *Zip10^d/d^* mice. (**A**) The representative images of calcium spike and zinc spark after IVF in mouse oocytes. Left side images showed calcium spike. Right side images showed zinc spark. The oocytes increased calcium ion and released zinc ion shortly after fertilization. The white dotted circles indicate the positions of oocytes. Successful fertilization was confirmed by simultaneously monitoring intracellular calcium oscillations with Calbryte 590 AM and excellular zinc ions with FluoZin-3 every 4 s. Capacitated frozen-thawed sperm was added to MII at 2 min after imaging start. Orange line showed calcium ion and dark blue line showed zinc ion. Intracellular calcium increases immediately before a zinc spark. Scale bars denote 20 μm. (**B**) The representative images of an MII egg activated with 5 μM ionomycin followed by monitoring of intracellular calcium oscillations with Calbryte 590 AM and extracellular zinc using 20 μM FluoZin-3. The ionomycin was added to MII at 2 min after imaging start. Orange line showed calcium ion and dark blue line showed zinc ion. Intracellular calcium increases immediately before a zinc spark. Scale bars denote 20 μm.

### *Zip10^d/d^* mouse oocytes can be fertilized but were unlikely to develop to the **blastocysts.**

*Zip6^d/d^* and *Zip10^d/d^* oocytes then were used for IVF, and we examined the rates of fertilization, polyspermy, and embryo development. Successful fertilization was confirmed at 6 h after IVF by the presence of pronuclei (Fig. 4A and B). *Zip10^d/d^* oocytes were fertilized at rates like those observed for control oocytes (Fig. 4A). Consistent with this, the zona reaction triggered by fertilization and examined using a ZP2 antibody was similar in *Zip10^d/d^* and *Zip10^f/f^* oocytes (Fig. 4D). We also examined the localization of ovastacin, whose expression in the cortex and loss post-fertilization was comparable between the two groups (Fig. 4E). As for the expression of JUNO, it had the same expression than between null and control oocytes (Fig. S4) and the temporal dynamics of its disappearance from the cortex after fertilization was similar for both *Zip10^f/f^*and *Zip10^d/d^* groups (Fig. 4F).

**Fig. 4.**
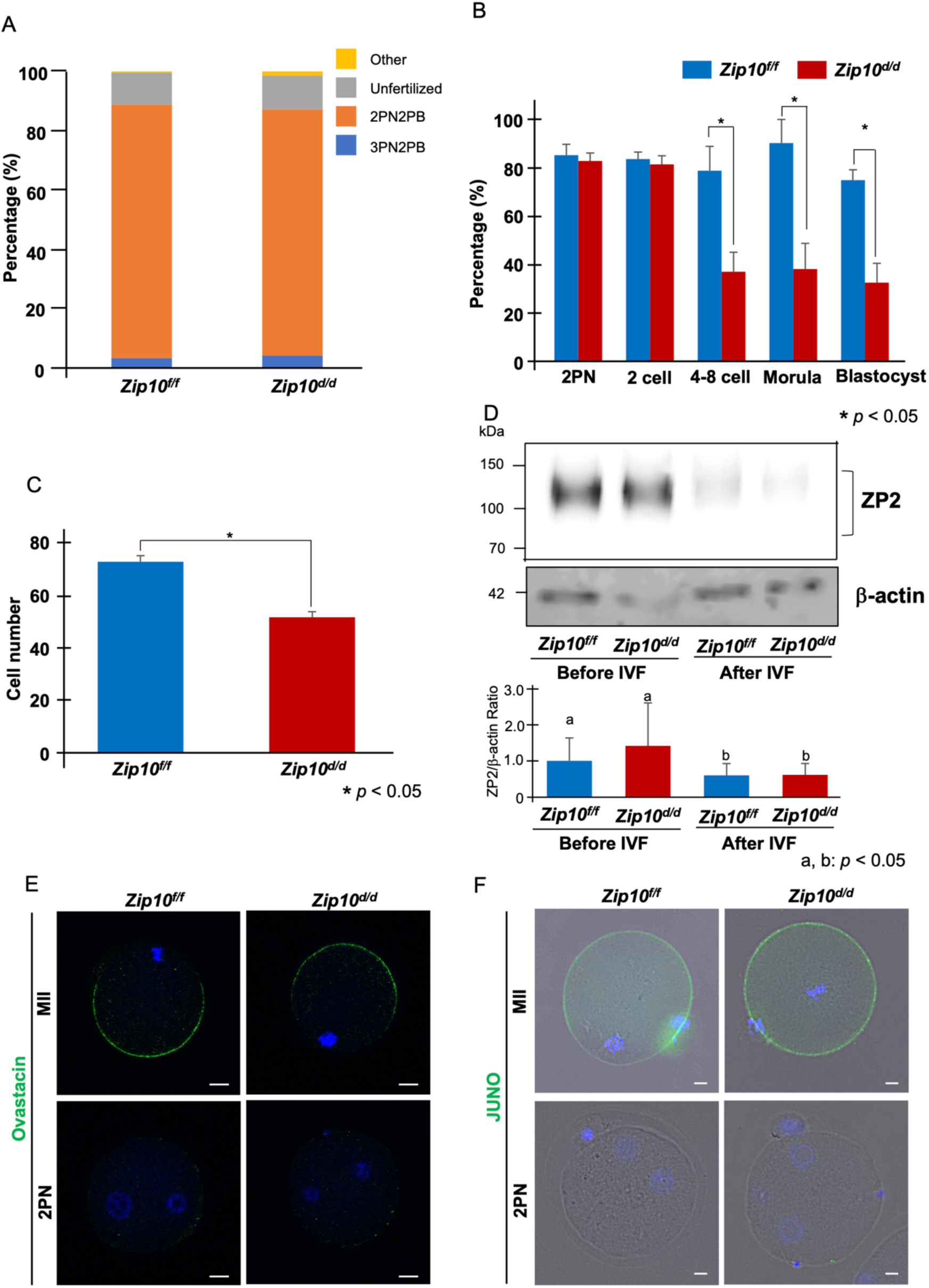
Presence of a mechanism to prevent multisperm fertilization in Zip10 cKO mice. (**A**) The percentages of oocytes with each number of PN at 6 h after insemination. Yellow region showed other including degeneration, degression and fragmentation. Gray region showed unfertilization, namely MII oocytes. Orange showed 2PN2PB, namely embryo possessed one female and male pronucleous (2PN) and second polar body (2PB). Blue region showed multisperm fertilization (3PN2PB). (**B**) The percentage of fertilized oocytes and developmental embryos. Data represent the average ± SE of the experiments. The embryo development was observed at 6 (2PN) 24 (2 cell), 48 (4-8 cell), 72 (Morula), and 96 (Blastocyst) hours after IVF. The oocytes used for IVF were calculated as the parameter for the fertilization rate and the rate of embryo development. These experiments were repeated at least thrice. Statistical differences were calculated according to the chi-square test. Different letters represent significant differences (*p* < 0.05). (**C**) The cell number of blastocyst derived from IVF. Blastocysts were fixed, immunostained, and physically flattened between a slide and coverslip. Photographs represent a single plane of focus. Nuclei are indicated by DAPI staining. The count used inverted fluorescence microscope. These experiments were repeated three times, and each group counted total 46 embryos. Statistical differences were calculated according to the student’s *t*-test (*p* < 0.05; significant difference). (**D**) Western blot of oocytes from *Zip10^f/f^* and *Zip10^d/d^* mice at 0 or 6 h after insemination using rat anti-ZP2 antibody. Intact ZP2 and the cleaved C-terminal fragment of ZP2 measured 120-130 kD and undetected, respectively. Expression level of β-actin serves as a protein loading control and quantified the expression level of ZP2. Molecular mass is indicated at the left. Statistical differences were calculated according to the one-way ANOVA. Different letters represent significant differences (*p* < 0.05). (**E**) MII oocytes and 2PN embryos from *Zip10^f/f^* and *Zip10^d/d^* mice were imaged by confocal microscopy after staining with rabbit anti-ovastacin (green). Chromosomes were stained with DAPI (blue). The scale bar represents 10 μm. (**F**) MII oocytes and 2PN embryos from *Zip10^f/f^* and *Zip10^d/d^* mice were imaged by BZ-X700 microscopy after staining with rat anti-mouse FR4 (JUNO; green). Chromosomes were stained with DAPI (blue). The scale bar represents 10 μm.

We next examined the development to the blastocyst stage, and whereas *Zip10^f/f^* zygotes developed at the expected rates, approximately 78.9% (2 cell), 90.3 (4-8 cell) and 75.0% (blastocyst), a smaller fraction of *Zip10^d/d^*zygotes did 37.2% (2 cell) and 38.3% (4-8 cell) and only 32.7% reached blastocyst stage (*p* < 0.05; Fig. 4B). As shown in Fig. 4C, the total cell numbers of blastocysts were also lower for *Zip10^d/d^*embryos (51.6 ± 2.1 cells) than for those of *Zip10^f/f^* ones (72.7 ± 2.1 cells, *p* < 0.05). *Zip6^d/d^* oocytes did not display any alterations in the rate of fertilization or development (Fig. S2C and D; *p* > 0.05).

## Discussion

The zinc sparks associated with fertilization were first discovered in the mouse (Kim et al., 2011). Similar observations followed in other mammalian species and amphibians (Que et al., 2019; Duncan et al., 2016; Seeler et al., 2021). However, the underlying mechanism(s) and biological role of zinc sparks are not elucidated. In this study, we have examined the role of the zinc transporters ZIP6 and ZIP10 using conditionally gene deficient mice, and queries their contribution to the zinc sparks of fertilization.

First, we confirmed the expression of zinc transporter in mouse oocytes. Our results are consistent with previous studies (Kong et al., 2014; Chen et al., 2023). ZIPs, including ZIP6 and ZIP10, are transmembrane proteins (Eide, 2004; Kambe et al., 2004; Lu and Fu, 2007; Schmitt-Ulms et al., 2009). In primordial follicles, the ooplasm staining of ZIP10 we anticipate corresponds to ooplasmic vesicular sites. ZIP10 expression shifted to the plasma membrane in primary and antral follicle phase. A similar localization shift of ZIP10 to the oocyte surface was reported during oocyte maturation, which is much later than reported here by. Furthermore, in that study, ZIP10 was detected in the nuclear/nucleolar positions of oocytes of all follicular stages (Chen et al., 2023), which we did not observe. On the other hand, ZIP6 was expressed at the nuclear/nucleolar regions and in granulosa cells. This localization of ZIP6 was consistent with that in a previous study (Che et al., 2023). However, our results failed to notice that ZIP6 shifts to the at plasma membrane in antral follicle oocytes (Kong et al., 2014; Chen et al., 2023). The results indicate that ZIP6 may be fulfilling a distinct function within the oocytes compared to ZIP10.

To assess the role of these transporters in mice, we generated oocyte-specific *Zip6 or Zip10* knockout mice (Fig. S1; *Zip6^d/d^* and *Zip10^d/d^*, respectively). Previous reports noted that the disruption of these transporters using specific morpholinos or incubation in function-blocking antibodies induced the change of intracellular labile zinc quota into mice oocytes (Kong et al., 2014). The amount of labile zinc ions in those mouse oocytes was measured using Fluozin-3AM, zinc indicator. The amount of labile zinc ions in *Zip10^d/d^* oocytes was significantly lower than in the *Zip10^f/f^* (Fig. 2D). We failed to observe any effect on zinc levels of in *Zip6^d/d^* oocytes (Fig. S2B). The results indicate that ZIP10 is mostly responsible for the uptake of zinc ions in mouse oocytes.

Zinc insufficient GV oocytes do not maintain the meiotic arrest at PI (Kong et al., 2012). This is suggested to happen by premature activation of the MOS-MAPK pathway in the presence of low zinc (Kong et al., 2012). In addition, low ooplasmic zinc accelerates meiotic progression that contributes to the extrusion of large polar bodies (Bernhardt et al., 2011). The early mitotic inhibitor 2 (EMI2), a zinc-binding APC/C proteasome inhibitor, is also an essential component of the cytoplasmic factor (CSF) that initiates entry into MII phase. EMI2 is a zinc binding protein, and when zinc is reduced, the activity of the APC/C proteasome is stimulated, increasing the degradation of CCNB1, reduced MPF activity leading to early release of meiosis arrest (Bernhardt et al., 2011; Bernhardt et al., 2012; Suzuki et al., 2010; Shoji et al., 2014; Ohe et al., 2010). Previous studies reported that targeted disruption of *Zip10* using morpholino injections and function-inhibiting antibodies during meiotic maturation perturbed meiosis progression and resulted in cell cycle arrest at the telophase I-like state (Kong et al., 2014). Based on these findings we hypothesized that the reduced state of labile zinc ions in *Zip10^d/d^* mouse oocytes may resemble the state of zinc-deficient oocytes. We investigated the maturation progression and spindle organization in oocyte matured in vitro from the GV to MII stage. We failed to observe any differences (Fig. 2A, B and C). We also did not observed abnormalities in oocytes from *Zip6^f/f^* mice (Fig. S2A; *p* > 0.05). We speculate that the disparate outcomes observed in previous studies may be attributed to the presence of trace amounts of labile zinc ions in media or in the oocytes.

Surprisingly, the amount of labile zinc ions in *Zip10^d/d^*2PN zygote was higher than in *Zip10^f/f^* and despite some differences in fluorescence intensity between *Zip10^d/d^* GV, MII and 2PN zygote, the labile zinc concentrations were not significantly different between these groups (Fig. 2D). Therefore, we monitored zinc sparks and calcium oscillations in the *Zip10*-KO oocytes. Several studies reported the important of IP_3_Rs in mammalian oocytes (Fissore et al., 1999; Parrington et al., 1998), which is essential for egg activation because its inhibition precludes Ca^2+^ oscillations (Miyazaki and Ito, 2006; Miyazaki et al., 1992; Xu et al., 2003). It has been reported that a putative Zinc-finger motif in a helical linker (LNK) domain near the C-terminus of *IP_3_R1* plays a role in *IP_3_R1* function (Fan et al., 2015; Paknejad and Hite, 2018). Recently, Akizawa *et al*., reported that both deficient and excessive zinc ions compromise *IP_3_R1* sensitivity, diminishing and terminating calcium oscillations (Akizawa et al., 2023). Our results showed that a single zinc spark occurs immediately after first calcium rise of oscillations in *Zip10^f/f^* oocytes, as reported by Kim *et al* (Kim et al., 2011) (Fig. 3A), and similar results are obtained in *Zip6^d/d^* and *Zip6^f/f^*mouse oocytes (Fig. S3). However, in *Zip10^d/d^* oocytes, despite the presence calcium elevations, zinc sparks failed to occur. Similarly, *Zip10^d/d^* oocytes activated with ionomycin (artificial oocyte activation) did not show a zinc spark (Fig. 3B). In this study, calcium oscillations occurred in oocytes with low zinc state. In addition, there was no difference in the amplitude frequency of calcium ions in oocytes of both groups within the observation time. Akizawa *et al*., produced zinc deficiency state with TPEN, a strong chelator of intracellular labile zinc ions (Akizawa et al., 2023). On the other hand, we did not completely remove intracellular labile zinc ions from mouse oocytes. We speculate that zinc ions existed in *Zip10^d/d^*mouse oocytes induce Ca^2+^ release without compromising *IP_3_R1* sensitivity. However, this study demonstrated that the accumulation of intracellular zinc ions mediated by ZIP10 is essential for inducing zinc sparks in mouse oocytes.

The total cellular zinc ions content required for meiosis in the oocyte substantially increases (a 50% increase) from the prophase I arrest to the arrest at metaphase of meiosis II (Kim et al., 2010; Bernhardt et al., 2011). The zinc is stored in undefined cortical granules, and an average of 10^6^ zinc atoms are released from these vesicles at the time of fertilization (exocytosis) (Kim et al., 2011; Que et al., 2015; Kong et al., 2014; Que et al., 2019; Jo et al., 2019; Lee et al., 2020). There are several possibilities that might explain why *Zip10* conditional KO mice have low zinc sparks. The most logical is that these oocytes experience reduced zinc loading of cortical of zinc vesicles, leading to the absence of sparks. Our results suggest that in oocytes, zinc sparks and exocytosis require adequate concentrations of labile zinc ions in the cortical secretory vesicles, which is not accomplished when ZIP10 is not present.

Zinc sparks have been reported to contribute to the rapid and permanent block mechanisms to prevent polyspermy. The former is achieved by zinc in the extracellular milieu decelerating the forward motility of sperm (Tokuhiro and Dean, 2018). Secondly, the release of labile pools of zinc at the same time that the zinc metalloendopeptidase ovastacin is released with the cortical granules (Tokuhiro and Dean, 2018; Burkart et al., 2012; Schmitz et al., 2021), cause ZP2 protein cleavage inducing a complete, although belated block to polyspermy (Que et al., 2017; Aonuma et al., 1981; Tokuhiro and Dean, 2018). However, the physiological significance of the zinc sparks’ contributions to polyspermy have not been thoroughly tested, and *Zip10^d/d^*mice, which are almost devoid of them, offer a great model. Our results showing normal rates of monospermic fertilization in *Zip10^d/d^ oocyte* suggest that contributions of Zinc to polyspermy control is negligible (Fig. 4A). Our results also show that the ovastacin undergoes normal release in *Zip10^f/f^* oocytes after fertilization (Fig. 4E and 4D), confirming previous studies of a minor role of the ZP2 to polyspermy. We observed the disappearance of JUNO after fertilization (Bianchi et al., 2014) *Zip10^d/d^,* suggesting that gamete fusion took place (Fig. 4F). Together, these results support the view that zinc spark is not directly involved in the polyspermy rejection mechanism.

Interestingly, there were no differences in the rates of fertilization (pronucleus formation), but development to the blastocyst stage was significantly reduced in *Zip10-*null embryos (Fig. 4B). Most of the embryos were arrested at the 2-cell stage and not developed beyond the 4-cell stage. Zinc insufficiency caused altered chromatin structure in the nuclei of blastomeres that displayed decreased global transcription, causing arrest in embryonic development (Kong et al., 2015). The abnormal zinc homeostasis, particularly during the one-cell stage, inhibits the activation of the embryonic genome that occurs around the two-cell stage in mice due to reduce the translational capacity through the inhibition of ribosomal RNA synthesis by RNA polymerase I (Chanfreau, 2013; Garner et al., 2021). Our results also show that *Zip10^d/d^* blastocysts had significantly lower cell numbers compared to the blastocysts from *Zip10^f/f^* mice (Fig. 4C). Our results suggest that Zip10^d/d^ mice oocytes can be fertilizable, but their developmental potential is decreased. The lack of ZIP10-mediated zinc influx during the folliculogenesis of *Zip10^d/d^* may compromise gene expression during these early stages of oocyte development. *Zip10^d/d^* mouse oocytes thus may have compromised developmental potential from the outset. Future studies should assess the transcriptomic or proteomic profile of *Zip10^d/d^* mouse oocytes. In conclusion, we elucidated that ZIP10 is required for zinc ion uptake into oocytes and the intracellular zinc ions regulated by ZIP10 was important for zinc sparks and progression of embryonic development. However, the role of ZIP6 remained uncertain. Additionally, the absence of mechanistic insight for zinc spark and the inability to distinguish between the developmental and fertilization stage roles of ZIP10 remain unresolved. These challenges necessitate further investigation. Currently, the many infertility patients exist in the world (the survey of WHO). It is reported that approximately half of the adult females are deficient in zinc in their serum (Prasad, 1996; Yokokawa et al., 2020). Zinc has been reported to play essential roles in many physiological functions including reproductive functions (Suzuki, M., et al, 2021, MacDonald RS, 2000, Jackson KA, et al, 2008, Bernhardt ML, et al, 2012, Suzuki, T., et al, 2010, Ohe M, et al, 2010). The findings will help elucidate the role of zinc homeostasis in the fields of fertilization/embryogenesis. Furthermore, the development of zinc-focused assisted reproductive technologies and fertilization/embryogenesis media could contribute to improve the developmental potential of oocytes and embryos in other mammalian species.

## Materials and Methods

All chemicals and reagents were purchased from Sigma-Aldrich (St. Louis, MO, USA) unless otherwise stated.

### Animals

Animals were housed in the barrier facility at Azabu University. The deletion of the ZIP10 gene in the whole body is known to result in embryonic lethality (He et al., 2023). The conditional knockout mice were generated by Cre-loxP system. The following mouse strains were used: *Zip10^flox/flox^* (*Zip10^f/f^*) mouse <B6;129-Slc39a10 <tm1.1Tfk>> (RBRC06221) (Hojyo et al., 2014; Miyai et al., 2014; Bin et al., 2017) was provided by the RIKEN BRC through the National BioResource Project of the MEXT/AMED, Japan. *Zip6^flox/flox^* (*Zip6^f/f^*) mouse <B6;129-Slc39a6> was generated by AdAMS. *Gdf9^Cre/+^* mice <Stock Tg (GDF9-iCre) 5092Coo/J, Stock No: 011062> were delivered from Jackson Laboratory (Lan Z, et al, 2004). Crlj: C57BL/6J female mice (6-8 weeks old) and Crlj: C57BL/6J male mice (8-12 weeks old) were purchased from Charles River Laboratories Japan (Yokohama, Japan). The mice were housed under controlled lighting conditions (daily light period, 0600-1800 h). The study was approved by the Animal Experimentation Committees of Azabu University and were performed in accordance with the committees’ guidelines (200318-13 and 230309-1).

Oocyte-specific gene knockout (*Zip10^f/f^, Gdf9^Cre/+^*; *Zip10^d/d^*) mice were generated by crossing *Gdf9^Cre/+^* male mice with *Zip10^f/f^* female mice (Fig. S1A-E). Similarly, *Zip6^d/d^* mice were generated (Fig. S1A-E).

### In situ hybridization (ISH)

ISH was performed as previously described with some modifications (Namiki et al., 2023). Briefly, fixed ovaries were paraffin-embedded and paraffin sections (6 µm) were mounted on MAS-coated slides (Matsunami Glass Industries, Osaka, Japan) under RNase-free conditions. Sense or antisense digoxigenin (DIG)-labeled RNA probes for *Slc39a10* (*Zip10*) were purchased from Genostaff. The sections were deparaffinized, rehydrated, and post-fixed in 10% neutral buffered formalin (NBF) for 30 min at 37°C, followed by the treatment with 0.2% hydrogen chloride and 5 µg/ml proteinase K (FUJIFILM Wako Pure Chemical, Osaka, Japan) for 10 min at 37°C, respectively. Hybridization was performed with DIG-labeled probes (250 ng/ml) in a humidified chamber at 60°C overnight. The slides were washed after hybridization, then treated with blocking reagent (Genostaff) for 15 min and alkaline phosphatase-conjugated anti-DIG antibody (1:2,000; Roche Diagnostics, Basel, Switzerland) for 1 h at room temperature. The signals were detected by 4-nitro-blue tetrazolium/5-bromo-4-chloro-3-indolyl phosphate (NBT/BCIP, Roche Diagnostics) in a humidified container for 12 h at 4°C. The sections were counterstained with Kernechtrot solution (Muto Pure Chemicals, Tokyo, Japan). Signals detected by the sense probe were used as a control for background levels.

### Immunofluorescent staining (IF) for ovary

IF was performed as previously described with some modifications (Namiki et al., 2023). Fixed ovaries were paraffin-embedded and the sections (6 µm) were deparaffinized, hydrated and conducted for antigen retrieval by autoclaving in 10 mM sodium citrate buffer (pH = 6.0) for 5 min. The sections were further incubated on ice for 30 min. After blocking with Bloking One Histo (06349-64, NACALAI TESQUE Inc, Kyoto, Japan) for 1 h, the slides were incubated with primary antibody for Rabbit anti-mouse-Zip10 (1:200; (Miyai et al., 2014)), Rabbit anti-SLC39A6 (1:200, HPA042377, Sigma), Rabbit anti-Foxl2 (1:300; (Cocquet et al., 2002; Polanco et al., 2010)), Rat anti-ZP2 (1:100, sc-32752, Santa Crus Biotechnology, Dallas, Texas, USA) in Can Get Signal immunostain (TOYOBO, Tokyo, Japan) overnight at 4°C. The slides were incubated with Alexa Fluor 488 donkey anti-rabbit IgG (H+L), Alexa Fluor 594 donkey anti-rabbit IgG (H+L) or Alexa Fluor 488 donkey anti-rat IgG (H+L) conjugated secondary antibodies (Jackson Immuno Research Laboratories, West Grove, PA, USA) diluted 1:500 in Can Get Signal immunostain (TOYOBO) for 1 h, and mounted with ProLong Glass Antifade Mountant with NucBlue Stain (P36981, Thermo Fisher Scientific, Waltham, MA, USA). Micrographs were captured by BZ-X700 microscopy (Keyence, Osaka, Japan).

### Oocyte preparation

Germinal vesicle (GV) oocytes were collected in the manner described in our previous study (Ito et al., 2008) with some modifications. Ovaries were collected from female mice 48 h after intraperitoneally injection with 5 IU equine chorionic gonadotropin (eCG) (PMS; Nippon Zenyaku Kogyo, Fukushima, Japan). They were placed in a 35 mm culture dish containing MEMα (no nucleosides, powder; Gibco/Thermo Fisher Scientific, Tokyo, Japan) medium with 26 mM NaHCO_3_, 75 mg/l penicillin, 50 mg/l streptomycin sulfate, 5% (v/v) heat-treated fetal calf serum (FCS), and 10 ng/ml epidermal growth factor (EGF). Cumulus-oocyte complexes (COCs) were released from the antral follicles by gentle puncturing with a needle.

To obtain metaphase II (MII) oocytes, the mice were intraperitoneally injected with 5 IU eCG followed by injection with 5 IU human chorionic gonadotropin (hCG) (Gonatropin; ASKA Pharmaceutical, Tokyo, Japan) at 48 h later. COCs were collected from the oviductal ampulla 14-16 h after hCG injection.

These oocytes were used in the following experiments.

### Count of ovulated oocytes

COCs-MII were obtained by superovulation treatment. The cumulus cells were removed from the COCs-MII with hyaluronidase (1 mg/ml) and gentle pipetting. All collected oocytes were counted and calculated as the number of ovulations.

### In vitro maturation (IVM) of GV oocytes

IVM were conducted in the manner described in our previous study (Kamoshita et al., 2021) with some modifications. Ovaries were placed in a 35 mm culture dish containing MEMα medium with 26 mM NaHCO_3_, 75 mg/l penicillin, 50 mg/l streptomycin sulfate, 5% (v/v) FCS, and 10 ng/ml EGF. COCs-GV were released from the antral follicles by gentle puncturing with a needle. The COCs-GV were washed 3 times and cultured in 500 µl of same medium in 4-well dish at 37°C in an atmosphere of 5% CO_2_ in air for 10, 12 and 14 h. After the culture, cumulus cells were removed from the COCs with hyaluronidase (1 mg/ml) and gentle pipetting. The extrusion of 1st polar body was evaluated at each time.

### IF for oocytes/preimplantation embryos

The α-tubulin and JUNO (PE anti-mouse FR4) were performed with some modification of the methods of Inoue *et al* (Inoue et al., 2017). The oocytes or embryos were fixed in 4% PFA for 30 min at room temperature, washed in PBS containing 1% polyvinyl alcohol (PBS/PVA). They were permeabilized by treatment of 0.5% Triton X-100 for 15 min and washed 2 times in 1% BSA/PBS/PVA followed by blocked for 20 min in same medium. They were incubated for overnight at 4° with primary antibodies to rabbit anti-α-tubulin (1:200; 11H10, cell Signaling Technology) or rat anti-mouse FR4 (JUNO; 1:250; 12A5 BioLegend), wash 3 times in 1% BSA/PBS/PVA. Primary antibodies were detected using Alexa Fluor 488 donkey anti-rabbit IgG (H+L) (1:250) or Alexa Fluor 488 donkey anti-rat IgG (H+L) (1:250) for 1 h at room temperature. After staining, all samples mounted in VECTASHIELD Mounting Medium with DAPI (H-1200; Vector Laboratories, California, USA), imaged using BZ-X700 microscopy (Keyence, Osaka, Japan).

The ovastacin staining was performed with some modifications of the methods of Burkart *et al* (Burkart et al., 2012). oocytes or embryos were fixed in 4% PFA for overnight at 4 °C, washed in PBS containing 0.3% polyvinylpyrrolidone (PVP), and then blocked in 0.3% BSA/0.1 M glycine (three times for 10 min) followed by permeabilization in 0.2% Triton X-100 for 15 min (Baibakov et al., 2007). Samples were then incubated for overnight at 4°C with Rabbit polyclonal anti-ovastacin (Burkart et al., 2012), 1:200, gifted antibody), washed with 0.3% PVP/0.1% Tween (three times for 10 min), and incubated for 1 h at room temperature with Alexa Fluor 488 anti-rabbit secondary antibody (1:500) followed by staining and mounting with VECTASHIELD Mounting Medium with DAPI. Samples were imaged using TCS SP5 II confocal microscope (Leica Microsystems, Wetzlar, Germany).

### In vitro fertilization (IVF)

IVF and sperm collection were conducted with some modification of the method described in our previous study (Kageyama et al., 2023). In brief, ovulated COCs-MII were preincubated for 1 h in 80 µl human tubal fluid (HTF) droplets supplemented with 1.25 mM reduced glutathione (GSH). Frozen-thawed sperm suspensions were suspended in 200 µl preincubation medium (HTF containing 0.4 mM methyl-β-cyclodextrin) and 0.1 mg/ml PVA, but without bovine serum albumin, and were incubated at 37 °C under 5% CO_2_ in humidified air for 1 h. At the time of insemination, preincubated spermatozoa were transferred into the droplets with oocytes at final concentrations of 2.0×10^6^ sperm/ml. After 6 h, oocytes were separated from spermatozoa and cumulus cells using a fine glass pipette and transferred into 50 µl KSOMaa medium. They were cultured at 37 °C under 5% CO_2_ in humidified air for approximately 24-96 h. The embryos were observed at 24, 48, 72, and 96h after IVF, and the number of 2cell, 4-8cell, morula, and blastocyst stage embryos were counted, respectively. The oocytes used for IVF were calculated as the parameter for the fertilization rate and the rate of embryo development. After IVF at 96 h, the blastocysts were fixed in 4% PFA for 30 min at room temperature. Following three times washed in PBS/PVA for 30 min each. Nuclear DNA was stained and mounted in VECTASHIELD Mounting Medium with DAPI. Cell numbers were determined by visually inspecting nuclei stained with DAPI using an inverted fluorescence microscope.

### Zinc measurements

FluoZin-3AM staining was conducted using the method described in our previous study (Kageyama et al., 2022). The obtained GV oocytes, cumulus cells were removed from the COCs with gentle pipetting. The obtained MII oocytes, cumulus cells were removed from the COCs with hyaluronidase (1 mg/ml) and gentle pipetting. Oocytes with a polar body were defined as MII. After IVF for 6h, two pronucleus stage embryos (2PN) were collected. GV, MII and 2PN were loaded in 50 µl medium that was suitable for each stage containing the amyl ester of the membrane permeant zinc-specific fluorophore, FluoZin-3AM (2 µM; F24195, Thermo Fisher Scientific, excitation 494 nm/emission 516 nm) for 1 h in humidified CO_2_ [5% (v/v) in air] at 37 °C followed by washing three times in medium and then observation with TCS SP5 II confocal microscope. FluoZin-3 has been extensively characterized for measurements of free intracellular zinc in live cells using microscopy and has an affinity constant (Kd) for zinc of 15 nM (Gee et al., 2002a; Gee et al., 2002b). Our previous study showed MII oocytes were treated with FluoZin-3 AM for 60 min, the change of fluorescence was confirmed in the cytoplasm of the oocytes and embryos, suggesting this treatment duration with FluoZin-3 AM is suitable for detection of zinc ions in oocytes and embryos(Kageyama et al., 2022). The pixel intensity per unit area after background subtraction was determined in GV, MII and 2PN within the circle (white circle) and ImageJ image processing software.

### Measurement of Ca^2+^ and Zinc spark during fertilization

Fresh or frozen-thawed sperm suspension were suspended in 200 µl preincubation medium. The 1 µl (fresh) or 20 µl (frozen-thawed) sperm suspension were placed in 40-80 µl HTF drop in pre-insemination dish about 20 min before insemination. The zona pellucida were punctured by piezo-electric pulses applied to four locations with a 15 µm injection needle. The treated MII oocytes were cultured in CZB containing CalbryteTM 590 AM (10 µM, #20700, AAT Bioquest, CA, USA, excitation 581 nm/emission 593 nm) for 10 min in humidified CO_2_ [5% (v/v) in air] at 37 °C. While the oocytes are incubated for 10 min, make a drop of 10 µl of PVA(-), Ca(-) Hepes-CZB (H-CZB) and pull out 7 µl of medium from drop with pipette man in the insemination dish. After 10 min, the oocytes are washed and transferred to insemination dish and the oocytes are attached to the dish. After attachment, an HTF containing 7 µl of BSA and a membrane-impermeable zinc-specific fluorophore, FluoZin-3 (20 µM, F24194, Thermo Fisher Scientific, excitation 494 nm/emission 516 nm), is added to the 3 µl drop containing the oocytes gently using a capillary. The about 1 µl sperm were sucked from the pre-insemination drop with the capillary under a stereomicroscope and place into the drop containing the oocytes, to start imaging. In case ionomycin treatment, after calcium labeling, Mll were allowed to settle in 45 µl PVA(-), Ca(-) H-CZB containing FluoZin-3 medium drop on dish, to start imaging. The 5 µl ionomycin (5 µM; #407950) were added to MII at 2 min after imaging start. Imaging was performed on confocal microscope using 488 nm (Zn^2+^) and 555 nm (Ca^2+^) excitation (Nikon solutions, Tokyo, Japan) for max 50 min every 4 seconds. Imaging analysis was performed by defining regions of interest (ROIs) and measuring fluorescence intensity over time using NIS-Elements (Nikon). The intracellular ROIs were drawn as the entire interior area of the cell. The extracellular ROIs were defined as a ring around the perimeter of cell. The ring thickness was conserved for all data analyses.

### Western blotting

Western blotting was carried out as described (Ito et al., 2010) with some modifications. Thirty MII oocytes or 2PN embryos were lysed in Laemmli sample buffer (Bio-Rad Laboratories, Tokyo, Japan) with 5% 2-mercaptoethanol. Samples were separated on 8% Bis-Tris gels by SDS-PAGE and transferred to PVDF membranes (Bio-Rad). The PVDF membranes were blocked in 10% skim milk (FUJIFILM Wako) in Tris-buffered saline (TBS) with 0.1% Tween-20 (Yoneyama yakuhin kogyo, Osaka, Japan) and probed with primary antibody to Rabbit anti-mouse-Zip10 (1:1,000; (Miyai et al., 2014)), Rabbit anti-SLC39A6 (1:1,000, HPA042377), rat anti-ZP2 (1:1,000) or monoclonal mouse anti-β-actin (1:5,000; A5316, Sigma-Aldrich) for overnight at 4°C. The membranes were incubated with secondary antibody; HRP-conjugated anti-rabbit IgG (1:5,000; Cell Signaling Technology, Danvers, MA, USA), HRP-conjugated anti-rat IgG (1:5000; Cell Signaling Technology) or HRP-conjugated anti-mouse IgG (1:5,000; Cell Signaling Technology) for 1 h at room temperature. After washing of the membranes, immunoreactive proteins were visualized using ECL^TM^ Western Blotting Analysis System (Cytiva Global Life Sciences Technologies, Tokyo, Japan), according to the manufacture’s recommendation. The membranes were exposed by ImageQuant LAS 4000. After exposure, the membranes were incubated for 30 min at 50 °C in the stripping buffer including 1.5 M Tris-HCl (pH 6.8), 10% SDS, 2-mercaptoethanol followed by extensive washing of the membranes. The membranes were probed with primary antibody to mouse anti-β-actin (1:5000; Sigma-Aldrich) for overnight at 4°C. The membranes were incubated with secondary antibody; FRP-conjugated anti-mouse IgG (1:5000; Cell Signaling) for 1 h at room temperature. The same procedure was followed below. The intensities of ZP2 bands were measured by quantitative analysis by densitometry using Image J.

## Statistical analysis

Values from three or more times were used for evaluation of statistical significance. Statistical analysis was performed using Statcel 3 software (OMS Ltd., Saitama, Japan). The fertility, the total numbers of oocytes collected from each mouse, the percentage of first polar extrusions, the fluorescence intensity of Fluozin-3 AM, the cell count of blastocysts and Quantification of JUNO expression were evaluated statistically by Student’s or welch’s *t*-test analysis. The rate of fertilization and embryo development was analyzed using chi square tests. The Quantification of ZP2 expression before and after fertilization was analyzed using one-way ANOVA. Values are shown as means ± S.E.M, and significant differences were considered at *p* values < 0.05.

## Acknowledgments and funding sources

We thank the members of the Laboratory of Animal Reproduction, School of Veterinary Medicine, Azabu University for technical help. *Zip10^flox/flox^* (*Zip10^f/f^*) mouse <B6;129-Slc39a10 <tm1.1Tfk>> (RBRC06221) was provided by the RIKEN BRC through the National BioResource Project of the MEXT/AMED, Japan. We also thank Dr. Austin J. Cooney (The University of Texas at Austin, Austin, United States) for providing GDF9-iCre mice. This research was supported by Grants-in-Aid for Scientific Research from the Japan Society for the Promotion of Science (JSPS) (KAKENHI, 21H02384 and 20H05373 to J.I., 22H04268, 24H02601 and 24K23214 to A.K.), JSPS KAKENHI Grant Number JP 15H04584 (AdAMS) to JI, the Sasakawa Scientific Research Grant from The Japan Science Society (2019-4044) to A.K. and Mishima Kaiun Memorial Foundation (A-135) to A.K. This work was supported by JSPS KAKENHI Grant Number JP 22H04922 (AdAMS). This study was also supported by Center for Diversity, Equity & Inclusion, Azabu University (A.K). This research was partially supported by the Center for Human and Animal Symbiosis Science, Azabu University and a research project grant awarded by the Azabu University Research Services Division to J.I.

## Author contributions

Conceptualization: A.K., J.I., Data curation: A.K., Methodology: N.O., T.W., A.O., Investigation: A.K., N.O., T.N., Y.K., Resources: M.O., Y.Y., T.F., Visualization: A.K., N.O., T.N., Supervision: A.O., T.F., R.A.F. J.I., N.K., Writing—original draft: A.K., Writing—review & editing: All the authors reviewed and edited the manuscript.

## Data and materials availability

All data are available in the main text or the supplementary materials.

## Movie legends

**Movie 1. Monitoring of intracellular calcium ions during fertilization of *Zip10^f/f^* oocytes.** The changes in Ca^2+^ were detected by CalbryteTM 590 AM every 4 second. *Zip10^f/f^* oocytes displayed the calcium oscillations following fertilization. The changes in zinc ions were also monitored simultaneously (Movie 2). The video was excerpted from maximum 50 min.

**Movie 2. Monitoring of extracellular zinc ions during fertilization of *Zip10^f/f^* oocytes.** The changes in Zn^2+^ were detected by FluoZin-3 every 4 second. *Zip10^f/f^* oocytes released zinc ions into the extracellular environment through a zinc spark followed the first calcium rise following fertilization. The changes in calcium ions were also monitored simultaneously (Movie 1). The video was excerpted from maximum 50 min.

**Movie 3. Monitoring of intracellular calcium ions during fertilization of *Zip10^d/d^* oocytes.** The changes in Ca^2+^ were detected by CalbryteTM 590 AM every 4 second. *Zip10^d/d^* oocytes displayed the calcium oscillations following fertilization. The changes in zinc ions were also monitored simultaneously (Movie 4). The video was excerpted from maximum 50 min.

**Movie 4. Monitoring of extracellular zinc ions during fertilization of *Zip10^d/d^*oocytes.** The changes in Zn^2+^ were detected by FluoZin-3 every 4 second. *Zip10^d/d^* oocytes did not release zinc ions into the extracellular environment followed the first calcium rise following fertilization. The changes in calcium ions were also monitored simultaneously (Movie 3). The video was excerpted from maximum 50 min.

**Movie 5. Monitoring of intracellular calcium ions during parthenogenesis of *Zip10^f/f^* oocytes.** The changes in Ca^2+^ were detected by CalbryteTM 590 AM every 4 second. *Zip10^f/f^* oocytes displayed a transient rise of calcium ions following artificial oocyte activation with ionomycin. The changes in zinc ions were also monitored simultaneously (Movie 6). The video was excerpted from maximum 50 min.

**Movie 6. Monitoring of extracellular zinc ions during parthenogenesis of *Zip10^f/f^* oocytes.** The changes in Zn^2+^ were detected by FluoZin-3 every 4 second. *Zip10^f/f^* oocytes released zinc ions into the extracellular environment through a zinc spark followed a transient calcium rise following artificial oocyte activation with ionomycin. The changes in calcium ions were also monitored simultaneously (Movie 5). The video was excerpted from maximum 50 min.

**Movie 7. Monitoring of intracellular calcium ions during parthenogenesis of *Zip10^d/d^* oocytes.** The changes in Ca^2+^ were detected by CalbryteTM 590 AM every 4 second. *Zip10^d/d^* oocytes displayed a transient rise of calcium ions following artificial oocyte activation with ionomycin. The changes in zinc ions were also monitored simultaneously (Movie 8). The video was excerpted from maximum 50 min.

**Movie 8. Monitoring of extracellular zinc ions during parthenogenesis of *Zip10^d/d^* oocytes.** The changes in Zn^2+^ were detected by FluoZin-3 every 4 second. *Zip10^d/d^* oocytes did not release zinc ions into the extracellular environment followed a transient calcium rise following artificial oocyte activation with ionomycin. The changes in calcium ions were also monitored simultaneously (Movie 5). The video was excerpted from maximum 50 min.

