## Supplementary Figure & Movie legends for "The oocyte zinc transporter *Slc39a10/Zip10* is a regulator of zinc sparks during fertilization in mice"

**Supplementary Information**

**Fig. S1.**

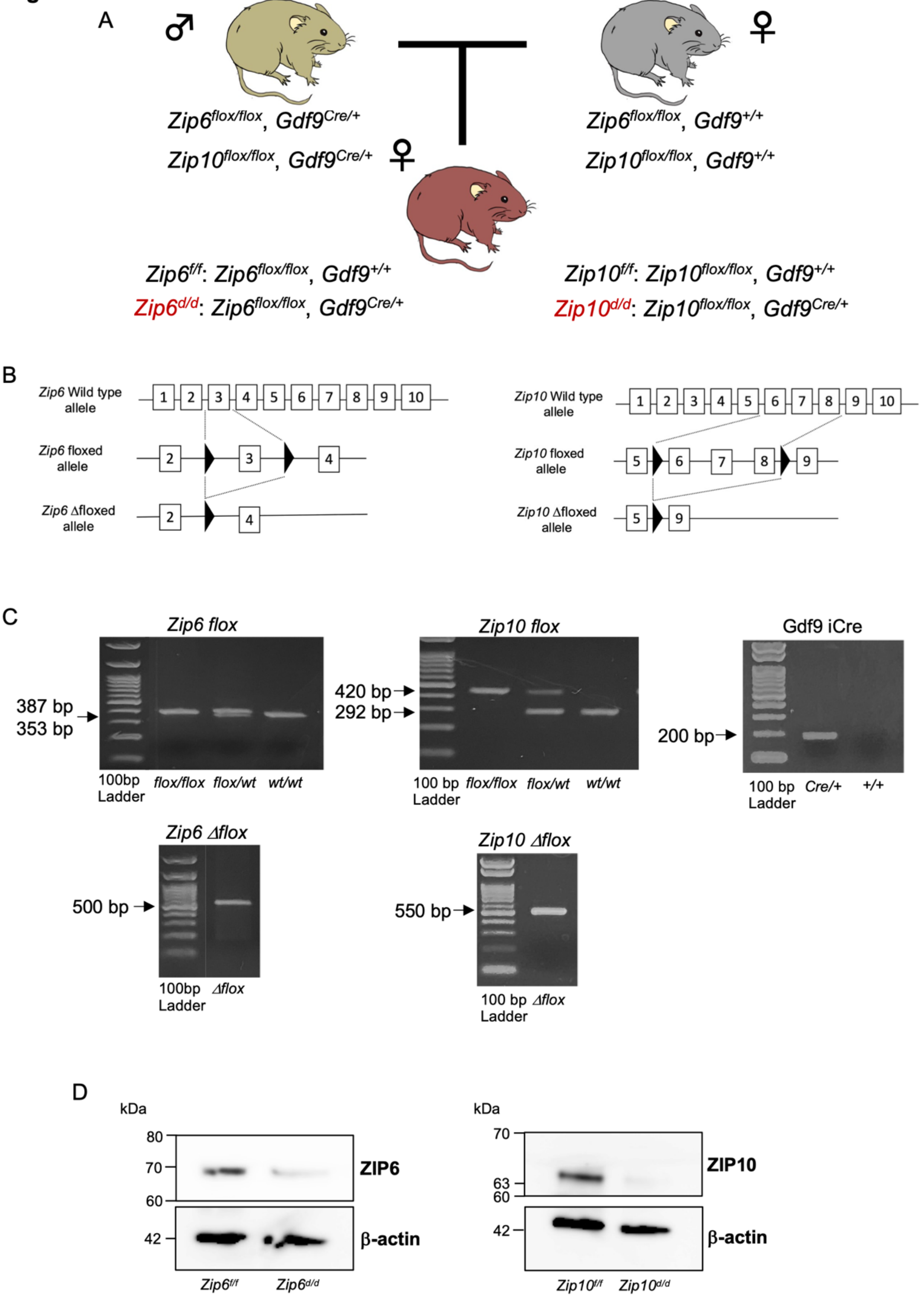

**Fig. S1 Generation of oocyte specific *Zip6* and *Zip10* conditional knockout mouse.** (**A**) Schematic of mating pattern of *Zip6* and *Zip10* conditional knockout mice. (**B**) Strategy used to develop floxed *Zip6* and *Zip10* allele. The mouse ZIP6 locus is shown at top. A loxP site (black triangle) sequence insert into a region before and after exon 3 of *Zip6* to produce the *Zip6* floxed allele with loxP sites flanking exons 3. In the presence of Cre recombinase there is further recombination resulting in the inactivated *Zip6* allele lacking exons 3 (*Zip6* ∆flox). The mouse ZIP10 locus is shown at top. A loxP site (black triangle) sequence insert into a region between exons 5 and 6, and a loxP site sequence between exons 8 and 9 of *Zip10* to produce the *Zip10* floxed allele with loxP sites flanking exons 6-8. In the presence of Cre recombinase there is further recombination resulting in the inactivated *Zip10* allele lacking exons 6-8 (*Zip10* ∆flox). (**C**) Genotyping of flox, ∆flox and iCre was performed by genomic PCR. To elucidate the roles of ZIP10 in the mouse oocytes, we crossed $Zip10^{flox/flox}$ ($Zip10^{f/f}$) mice with Growth Differentiation factor 9 (*Gdf9*)-Cre transgenic mice to generate $Zip10^{flox/flox}$ *Gdf9*-Cre ($Zip10^{d/d}$) mice. $Zip10^{f/f}$ mice were used as control. Similarly, we crossed $Zip6^{flox/flox}$ ($Zip6^{f/f}$) mice with *Gdf9*-Cre transgenic mice to generate $Zip6^{flox/flox}$ *Gdf9*-Cre ($Zip6^{d/d}$) mice. $Zip6^{f/f}$ mice were used as control. The tail tips of 2 weeks old mice were cut. The samples were lysed in 100 µl of DirectPCR Lysis Reagent (Viagen Biotech) with proteinase K (0.1 mg/ml) at 50°C for overnight. To inactivate proteinase K, the samples were incubated at over 80 °C for 1 h. The genotypes confirmed by polymerase chain reaction (PCR), as suggested by Miyai et al., (2014) (Miyai T, et al, 2014) and RIKEN BRC using CAAGGCCAGCCAAAATTCTA (A-F) and GCTTTCCTCCCATCCTGATT (A-R) to detect Wild-type (292 bp) and *Zip10* flox (420 bp), A-F and GTGGCATGCGTGGAAGTTAG (B-R) to detected *Zip10* flox null (550 bp) alleles. the genotyping of *Zip6* flox was confirmed using CCAGCATTGCCCTCTGTAAGAGTC (*Slc39a6*_F) and GCCTAAAGAAGATACACTGACACGACG (*Slc39a6*_R) to detect Wild-type (353 bp) and *Zip6* flox (387 bp), GGACCGGTTGCATAGAGGAG (*Slc39a6*_Null_F) and GAGGCAGGCGGATTTCTGAG (*Slc39a6*_Null_R) to detected *Zip6* flox null

(500 bp) alleles. Similarly, we detected *Gdf9* iCre/+ alleles (200 bp) using CAGGTTTTGGTGCACAGTCA (21218) and GGCATGCTTGAGGTCTGATTAC (25494) to suggest by Jackson Laboratory. MII oocytes were used in order to detect oocyte expression. (**D**) The expected molecular weight for ZIP6 is about 70 kDa and ZIP10 is about 63 kDa. Expression level of β-actin (42 kDa) served as a protein loading control. Molecular mass is indicated at the left.

Fig. S2.

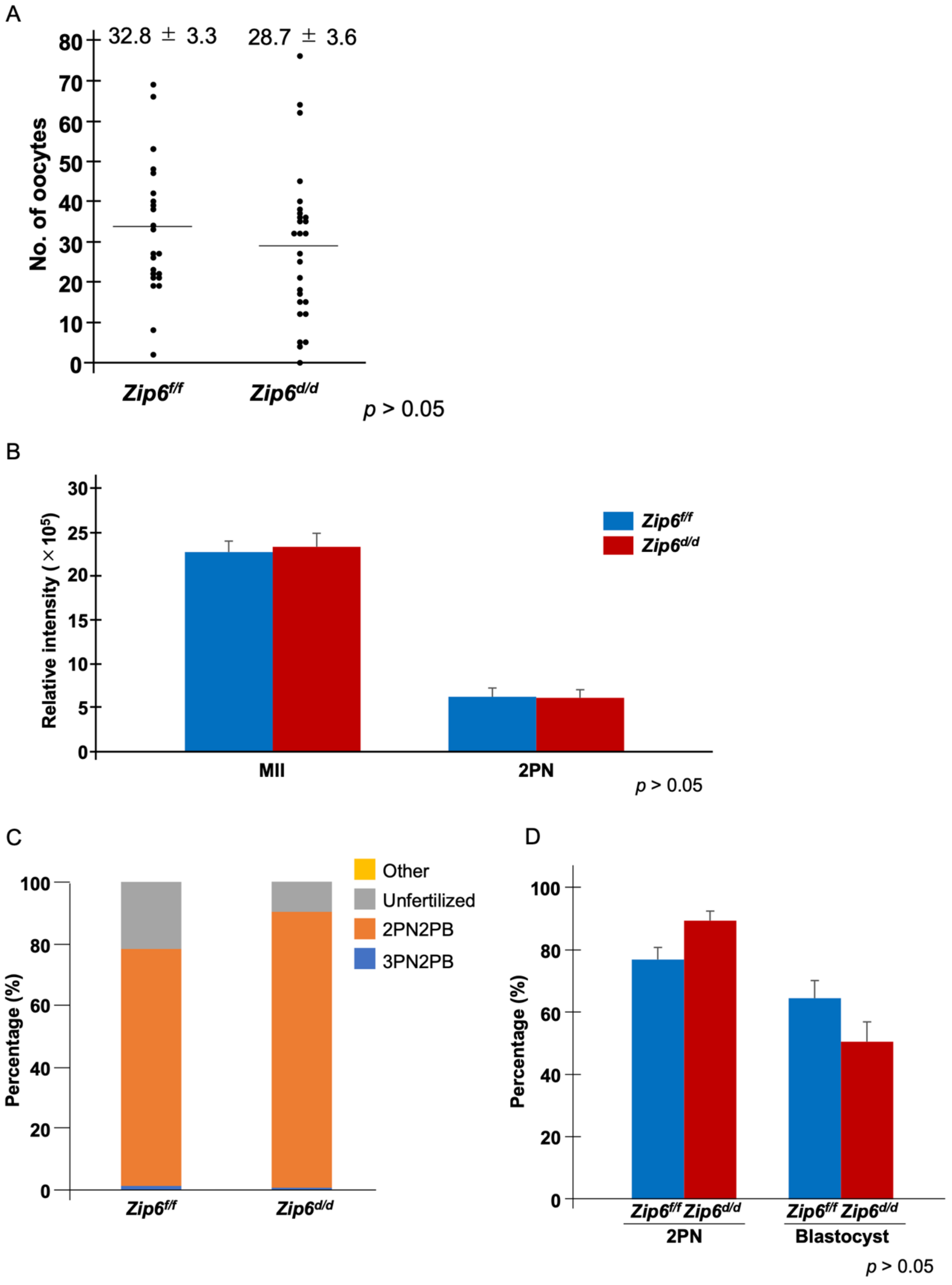

A
32.8 ± 3.3
28.7 ± 3.6
No. of oocytes
80
70
60
50
40
30
20
10
0
Zip6f/f
Zip6d/d
p > 0.05
B
Relative intensity (×10^5)
30
25
20
15
10
5
0
Zip6f/f
Zip6d/d
MII
2PN
p > 0.05
C
Other
Unfertilized
2PN2PB
3PN2PB
Percentage (%)
100
80
60
40
20
0
Zip6f/f
Zip6d/d
D
Percentage (%)
100
80
60
40
20
0
Zip6f/f Zip6d/d
2PN
Zip6f/f Zip6d/d
Blastocyst
p > 0.05

**Fig. S2 Number of collected oocytes, dynamics of labile zinc ion and percentage of fertilization in *Zip6*$^{d/d}$ mice.** (**A**) The results of average number of oocytes in each group. Data represents the average ± SEM. These experiments were repeated at least thrice. Statistical differences were calculated according to student's *t*-test ($p > 0.05$; no significant difference). (**B**) Comparison with the fluorescence intensity of intracellular labile zinc ion in MII and 2PN treated with 2 μM FluoZin-3AM for 1h. Data represent the average ± SE of the experiments. For each experiment, 10–20 oocytes/embryos were stained and used for the measurement in each stage of the experiment, and these experiments were repeated three times. Statistical differences were calculated according to student's *t*-test ($p > 0.05$; no significant difference). (**C**) The percentages of oocytes with each number of PN at 6 h after insemination. Yellow region showed other including degeneration, degression and fragmentation. Gray region showed as unfertilized, namely MII oocytes. Orange showed 2PN2PB, namely embryo possessed one female and male pronucleous (2PN) and second polar body (2PB). Blue region showed multisperm fertilization (3PN2PB). (**D**) The percentage of fertilized oocytes and developmental embryos. Data represent the average ± SE of the experiments. These experiments were repeated at least thrice. Statistical differences were calculated according to the chi-square test ($p > 0.05$).

**Fig. S3.**

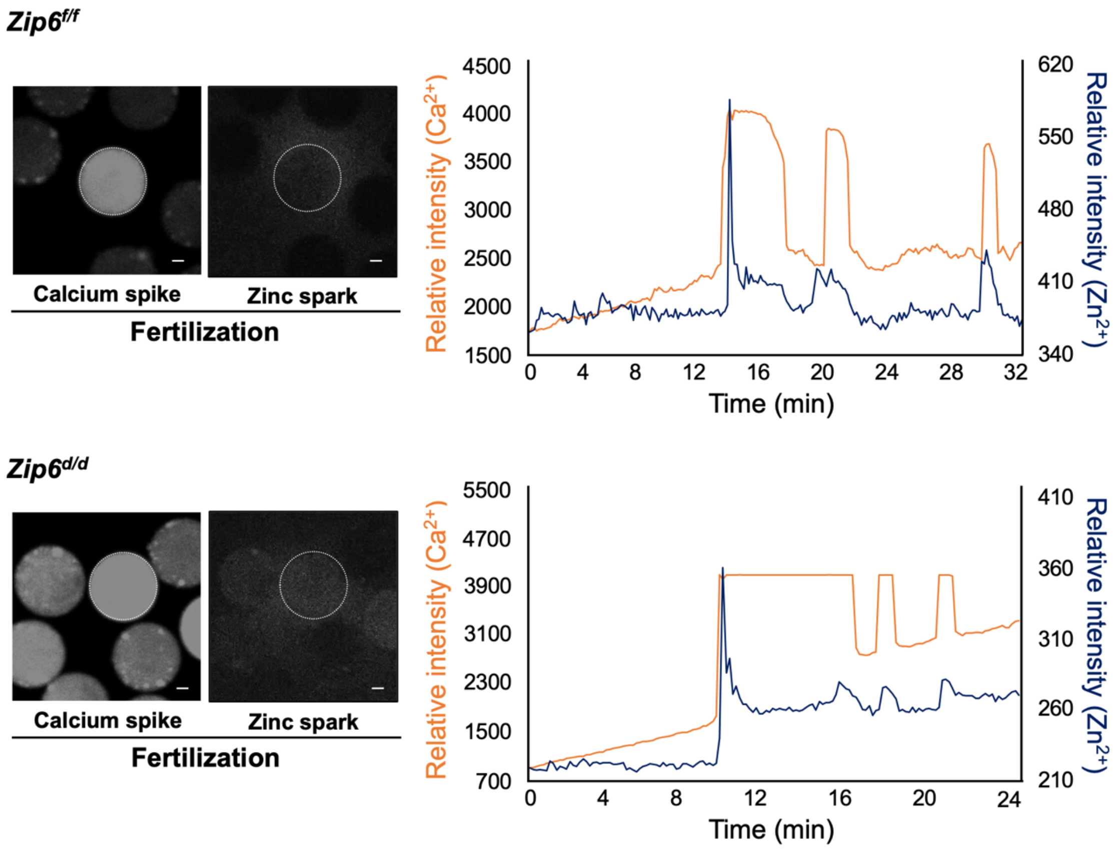


**Fig. S3 Measurement of calcium spike and zinc spark in *Zip6$^{d/d}$* mice.** The representative images of calcium spike and zinc spark after IVF in mouse oocytes. Left side images showed calcium spike. Right side images showed zinc spark. The oocytes increased calcium ion and released zinc ion shortly after fertilization. The white dotted circles indicate the positions of oocytes. Successful fertilization was confirmed by simultaneously monitoring intracellular calcium oscillations with Calbryte 590 AM and excellular zinc ions with FluoZin-3 every 4 s. Capacitated frozen-thawed sperm was added to MII at 2 min after imaging start. Orange line showed calcium ion and dark blue line showed zinc ion. Intracellular calcium increases immediately before a zinc spark. Scale bars denote 20 μm.

**Fig. S4.**

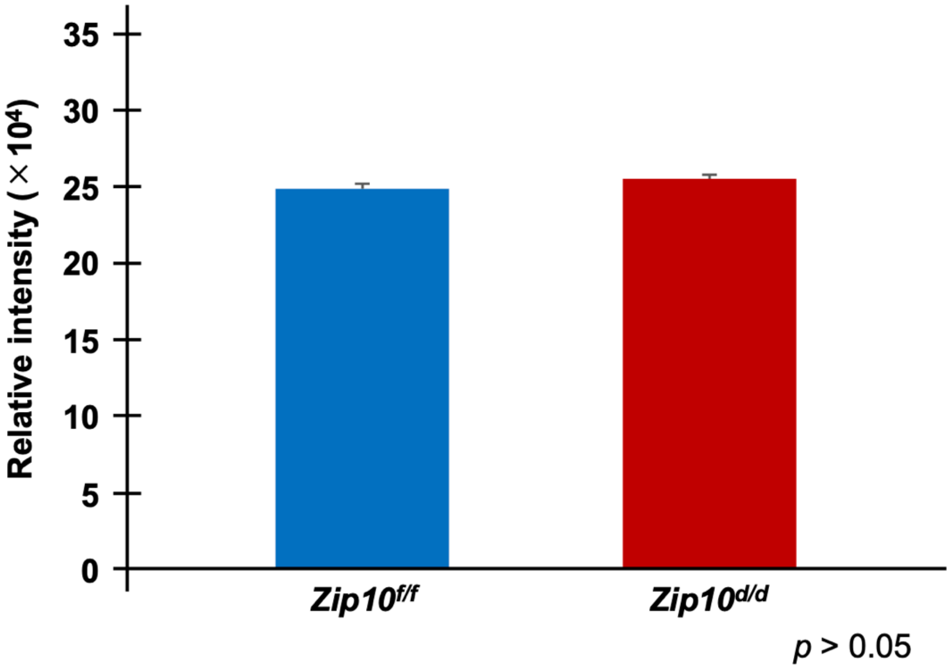


**Fig. S4 Comparison of JUNO expression in *Zip10$^{f/f}$* and *Zip10$^{d/d}$* mouse MII oocytes.** To measure JUNO-immunofluorescence intensity, oocytes images were selected as regions of interest (ROIs) and measured using ImageJ. Statistical differences were calculated according to student's *t*-test ($p > 0.05$; no significant difference).

**Supplementary Movie legends**

**Movie S1. Monitoring of intracellular calcium ions during fertilization of *Zip6^f/f^* oocytes.** The changes in $Ca^{2+}$ were detected by CalbryteTM 590 AM every 4 second. *Zip6^f/f^* oocytes displayed the calcium oscillations following fertilization. The changes in zinc ions were also monitored simultaneously (Movie S2). The video was excerpted from maximum 50 min.

**Movie S2. Monitoring of extracellular zinc ions during fertilization of *Zip6^f/f^* oocytes.** The changes in $Zn^{2+}$ were detected by FluoZin-3 every 4 second. *Zip6^f/f^* oocytes released zinc ions into the extracellular environment through a zinc spark followed the first calcium rise following fertilization. The changes in calcium ions were also monitored simultaneously (Movie S1). The video was excerpted from maximum 50 min.

**Movie S3. Monitoring of intracellular calcium ions during fertilization of *Zip6^d/d^* oocytes.** The changes in $Ca^{2+}$ were detected by CalbryteTM 590 AM every 4 second. *Zip6^d/d^* oocytes displayed the calcium oscillations following fertilization. The changes in zinc ions were also monitored simultaneously (Movie S4). The video was excerpted from maximum 50 min.

**Movie S4. Monitoring of extracellular zinc ions during fertilization of *Zip6^d/d^* oocytes.** The changes in $Zn^{2+}$ were detected by FluoZin-3 every 4 second. *Zip6^d/d^* oocytes released zinc ions into the extracellular environment through a zinc spark followed the first calcium rise following fertilization. The changes in calcium ions were also monitored simultaneously (Movie S3). The video was excerpted from maximum 50 min.
